# Impaired bone strength and bone microstructure in a novel early-onset osteoporotic rat model with a clinically relevant *PLS3* mutation

**DOI:** 10.1101/2022.06.29.498081

**Authors:** Jing Hu, Bingna Zhou, Xiaoyun Lin, Qian Zhang, Feifei Guan, Lei Sun, Jiayi Liu, Ou Wang, Yan Jiang, Weibo Xia, Xiaoping Xing, Mei Li

**Author notes:** Corresponding author to who reprint requests should be addressed: Mei Li. Department of Endocrinology, Key Laboratory of Endocrinology, National Health and Family Planning Commission, Peking Union Medical College Hospital, Chinese Academy of Medical Sciences and Peking Union Medical College, Shuaifuyuan No. 1, Dongcheng District, Beijing 100730, China.

## Abstract

Osteoporosis is a highly prevalent disorder with a strong genetic component. Recent studies identify *PLS3* as a novel bone regulator of bone metabolism and *PLS3* mutations lead to a rare monogenic early-onset osteoporosis. However, the mechanism of *PLS3* mutation leading to osteoporosis is unknown with no animal models carrying patient-derived *PLS3* mutations generated. The effective treatment strategies for *PLS3*-related have not been established. Here we have constructed a novel rat model with clinically relevant hemizygous E10-16del mutation of *PLS3* (*PLS3^E10-16del/0^*) that recapitulate the osteoporotic phenotypes with obviously thinner cortical thickness, significant decreases in yield load, maximum load, and breaking load of femora at 3, 6, 9 months compared to WT rats. Histomorphometric analysis indicates a significantly lower mineral apposition rate in *PLS3^E10-16del/0^*rats. Treatment with alendronate (ALN, 1.0 ug/kg per day) or teriparatide (TPTD, 40ug/kg five times weekly) for 8 weeks significantly improved bone mass and bone microarchitecture, and bone strength was significantly increased after TPTD treatment (P < 0.05). Thus, our results indicate the roles of *PLS3* in the regulation of bone microstructure and bone strength, providing a novel animal model for the study of early-onset osteoporosis. Alendronate and teriparatide treatment could be a potential treatment for early-onset osteoporosis induced by *PLS3* mutation.

## Introduction

Osteoporosis is the most common metabolic bone disorder and is characterized by low bone mineral density (BMD), bone microarchitecture deterioration, and increased predisposition for bone fractures. Osteoporosis is usually regarded as an age-related disease, however, idiopathic osteoporosis has also emerged in children and adolescents, which refers to significantly lower than expected bone mass manifesting in childhood with no identifiable etiology (O. Mäkitie & Zillikens, 2022). Recently, genetic variations have been found to be closely associated with diversities of bone mineral density (BMD), fracture risk and effects of anti-osteoporotic treatment (Ralston & Uitterlinden, 2010). A series of studies identified multiple heritable genetic variants that led to early-onset osteoporosis (EOOP).

In 2013, a new kind of severe EOOP was first described, which was caused by mutations in *PLS3* (van Dijk et al., 2013). *PLS3* (OMIM 300131) has 16 exons and spans approximately 90 kb, which is located on Xq23 and encodes a highly conserved protein plastin 3 (PLS3). PLS3 belongs to a family of actin-binding proteins, which is ubiquitously expressed in solid tissues and involved in the binding and bundling of actin filaments in the cytoskeleton, thus partaking in various cellular functions, such as cell migration and adhesion (Wolff et al., 2021). In bone, PLS3 is proposed to regulate cytoskeletal actin bundling, osteocyte function and their mechanosensory apparatus, and osteoclast function, as well as bone matrix mineralization (Pathak, Bravenboer, & Klein-Nulend, 2020; Wolff et al., 2021). A series of pedigree studies demonstrated that variants in *PLS3* led to significantly reduced BMD and early-onset recurrent fragility fractures (Costantini et al., 2018; Fahiminiya et al., 2014; Hu et al., 2020; Kampe et al., 2017; Kannu, Mahjoub, Babul-Hirji, Carter, & Harrington, 2017; Laine et al., 2015; Lv et al., 2017; van Dijk et al., 2013; Wang et al., 2020). Till now, 29 different mutations in *PLS3* have been reported to induce EOOP (Brlek et al., 2021; Cohen et al., 2022; Wolff et al., 2021). Due to its X-chromosomal inheritance, PLS3-induced osteoporosis has a more severe effect on males than females, although heterozygous carrier females also suffer from EOOP. However, the exact function of PLS3 in bone and the signal pathways in which it plays a role are still unknown.

So far, the animal models carrying patient-derived *PLS3* mutations have not been generated, which are valuable to unveil the pathogenesis of this ultra-rare osteoporosis induced by *PLS3* mutation, although there was a *PLS3* knock-out (KO) murine model (Neugebauer et al., 2018; Yorgan et al., 2020). Moreover, effective treatment strategies have not been established in EOOP related to *PLS3* mutation. Few patients with *PLS3*-associated EOOP received bisphosphonates or teriparatide treatment, while the efficacy was variable (Fratzl-Zelman et al., 2021; Hu et al., 2020; Lv et al., 2017; Valimaki et al., 2017; van Dijk et al., 2013). Triggered by a patient we previously reported with EOOP induced by a large fragment deletion in *PLS3* (Lv et al., 2017), we independently constructed a novel rat model with ubiquitous deletion of the exon 10-16 of *PLS3* (*PLS3^E10-16del/0^*), in order to explore the potential pathogenesis and treatment strategies of *PLS3*-related EOOP.

## Results

### The rats with hemizygous E10-16del mutation were successfully generated

A large fragment deletion of exon 10-16 in *PLS3* was introduced into the genome of rats, and a 9519-bp deletion from No. 84168bp to 93686bp (NC_051356.1) was confirmed by Sanger sequencing. Rats carrying mutant allele were born close to the expected Mendelian ratio (*PLS3^E10-16del^*^/+^: 28.6%, *PLS3^E10-16del^*^/0^: 28.5%, WT: 42.4%). The *PLS3^E10-16del^*^/0^ rats showed similar movement, locomotor activities, longitudinal growth and body weight compared to WT rats (Fig.1, Supplementary Fig.1).

**Fig. 1.**
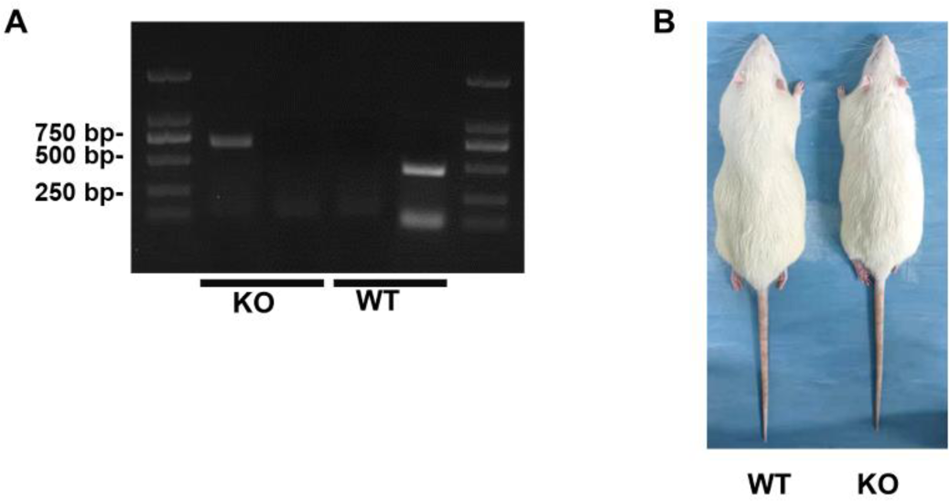
PCR genotyping and gross appearance in *PLS3^E10-16del/0^* and WT rats. Fig. 1A PCR genotyping of *PLS3^E10-16del/0^* rats. DNA extracted from tail snips was subjected to PCR using allele-specific primers for mutant and WT allele. Amplification of mutant samples resulted in one copy of the upper 653 bp fragment, while WT samples yield one copy of lower 450 bp fragment. Fig. 1B Gross appearance of 3-month-old male WT and *PLS3^E10-16del/0^* rats WT: wild type, KO: *PLS3^E10-16del/0^*

The biomechanical properties of *PLS3^E10-16del^*^/0^ rats were significantly impaired. Compared to age- and gender-matched WT group, *PLS3^E10-16del^*^/0^ rats exhibited a decrease by 24.8% in yield load (*P* < 0.001), 19.4% in maximum load (*P* < 0.01) and 25.3% in breaking load (*P* < 0.05) of femur in 3 months old, which continued to 6 and 9 months old (Fig. 2A, 2B). Notably, the stiffness (108.18±11.28 versus 501.20 ±84.97 N/mm, *P* < 0.001) and breaking load (155.99±39.92 versus 227.75±27.06 N, *P* < 0.01) of femur of 9-month old *PLS3^E10-16del^*^/0^ rats were pronouncedly lower than those of WT rats. Significantly reduced maximum load of vertebrae was also found in *PLS3^E10-16del^*^/0^ rats (Fig. 2C). Compared with WT rats, the average maximum load of the fifth lumbar vertebra of *PLS3^E10-16del^*^/0^ rats decreased by 28.5% (*P* < 0.01) and 43.6% (*P* < 0.001) at 3 and 6 months old. No significant difference was found in the stiffness, work-to-failure, and post-yield displacement between 3-month-old *PLS3^E10-16del^*^/0^ rats and WT rats (Fig. 2A, Supplementary Fig. 2A).

**Fig. 2.**
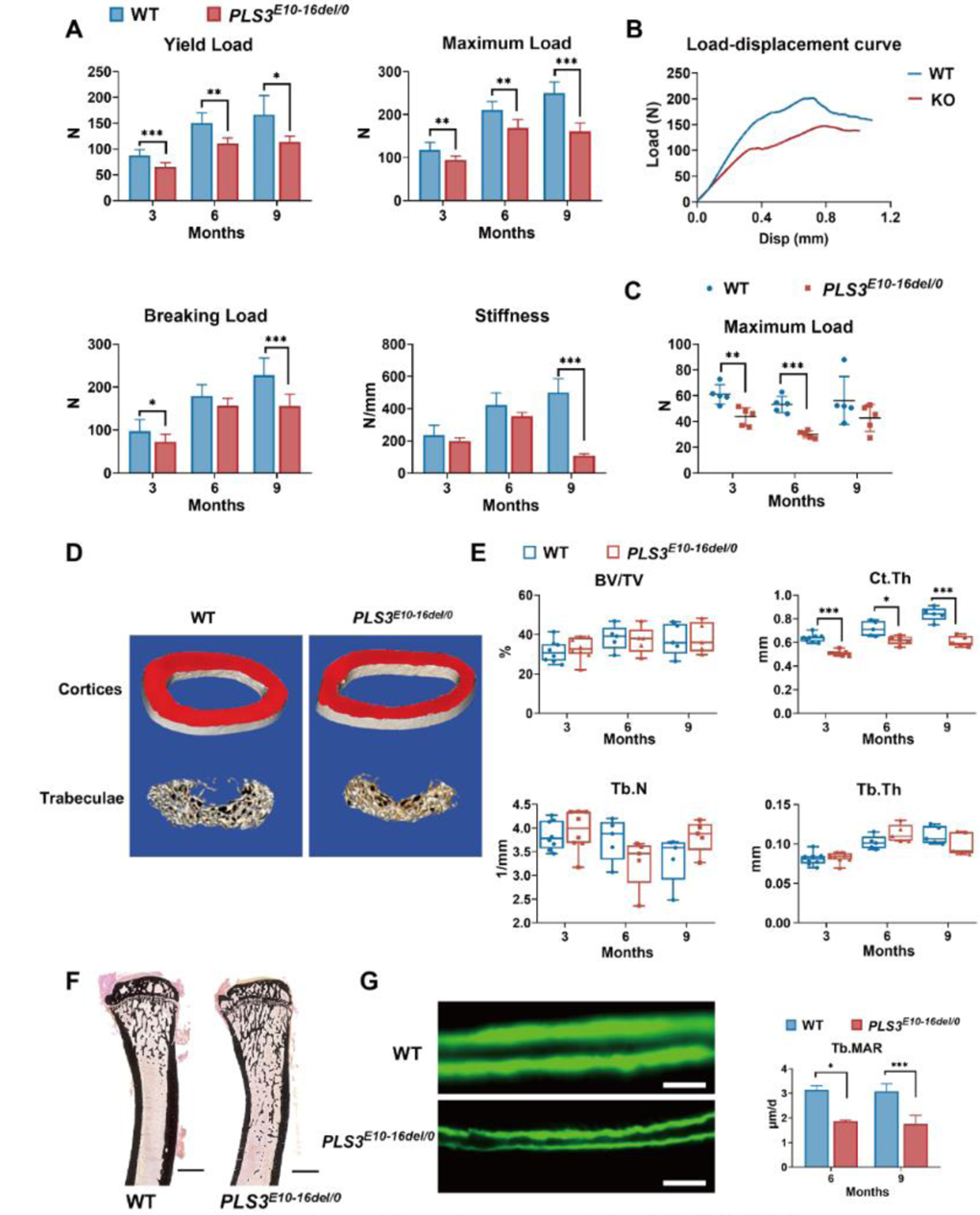
Bone strength, bone microstructure and bone formation activity of *PLS3^E10-16del/0^* rats. Fig. 2A Mechanical three-point bending tests of femora from *PLS3^E10-16del/0^* and WT rats (n= 5-8 per group) Fig. 2B Typical load-displacement curves of *PLS3^E10-16del/0^* and WT rats. Fig. 2C Indentation tests of L_5_ from *PLS3^E10-16del/0^* and WT rats \**P* < 0.05, ** *P* < 0.01, *** *P* < 0.001 versus WT groups Fig. 2D Three-dimensional reconstruction images of femurs from *PLS3^E10-16del/0^* and WT rats Fig. 2E Micro-CT assessment of the distal femurs from *PLS3^E10-16del/0^* and WT rats BV/TV: bone volume/tissue volume, Ct.Th: cortical thickness, Tb.Th: trabecular thickness, Tb.N: trabecular number Fig. 2F Representative von-Kossa-stained sections of tibia diaphysis of *PLS3^E10-16del/0^* and WT rats Scale bar = 2000 μm WT: wild-type Fig. 2G Typical images of unstained and uncalcified vertebra of *PLS3^E10-16del/0^* rats Scale bar = 10 μm, \**P* < 0.05 versus WT groups Tb.MAR: mineral apposition rate of lumbar trabeculae

The cortical bone microstructure of *PLS3^E10-16del^*^/0^ rats was deteriorated. Femoral cortical thickness of 3-, 6-, 9-month-old *PLS3^E10-16del^*^/0^ rats was 79.9% (*P* < 0.001), 86.3% (*P* < 0.05), 72.6% (*P* < 0.001) of age-matched WT rats, respectively. However, BV/TV, BS/BV, Tb.Th, Tb.N and Tb.Sp in femur were similar between *PLS3^E10-16del^*^/0^ rats and WT rats at all ages (Figure 2D, 2E, Supplementary Fig. 2B). No significant differences were found in %Tb.Ar, Tb.Th, Tb.N and Tb.Sp of lumbar vertebrae between the *PLS3^E10-16del^*^/0^ rats and WT rats (Supplementary Fig. 2C).

The mineral apposition rate (MAR) of trabecular bone (Tb.MAR) in lumbar vertebrae of 6- and 9-month-old *PLS3^E10-16del^*^/0^ rats and WT rats was significantly decreased (Fig. 2G), while no statistical changes were detected in MAR of endocortical (Ec.MAR) and periosteal surface (Ps.MAR) of tibia cortex (Supplementary Fig. 2D). The number of osteocytes, osteoclasts and osteoblasts were similar in *PLS3^E10-16del^*^/0^ rats and WT rats at all ages (Supplementary Fig. 2E). Serum levels of total ALP, β-CTX and calcium were similar in *PLS3^E10-16del^*^/0^ and WT rats. (Supplementary Fig. 2F)

### Effects of anti-osteoporotic treatment on *PLS3^E10-16del^*^/0^ rats

ALN and TPTD significantly improved bone microstructure of *PLS3^E10-16del^*^/0^ rats after 8-week treatment. Compared to vehicle (VEH) group, Tb.N and BV/TV values were increased by 38.4% and 38.7% in ALN group and by 35.9% and 29.3% in TPTD group, while Tb.Sp was decreased by 49.3% and 40.1% in ALN and TPTD group, respectively (all *P* < 0.05). Compared to VEH group, cancellous BMD of femur increased by 43.0% in ALN group (*P* < 0.01) and 33.3% in TPTD group (*P* < 0.01). Treatment of ALN and TPTD significantly increase 7.4% and 4.8% of cortical thickness of *PLS3^E10-16del^*^/0^ rats ( all *P* < 0.05) (Fig. 3A, 3B). A significant histomorphometric increase in %Tb.Ar (31.7%) and Tb.Th (60.3%) of L_4_ was observed in TPTD group (all *P* < 0.05), but not in ALN group (Fig.4A, 4B). Tibial Ec.MAR of TPTD group was higher than VEH group (*P* < 0.01), and rats in ALN group had the lowest Ec.MAR (2.75μm/d) in the tibia. (Fig. 4E). Ps.MAR of the tibia was similar among ALN, TPTD and VEH group (Supplementary Fig. 3C).

**Fig. 3.**
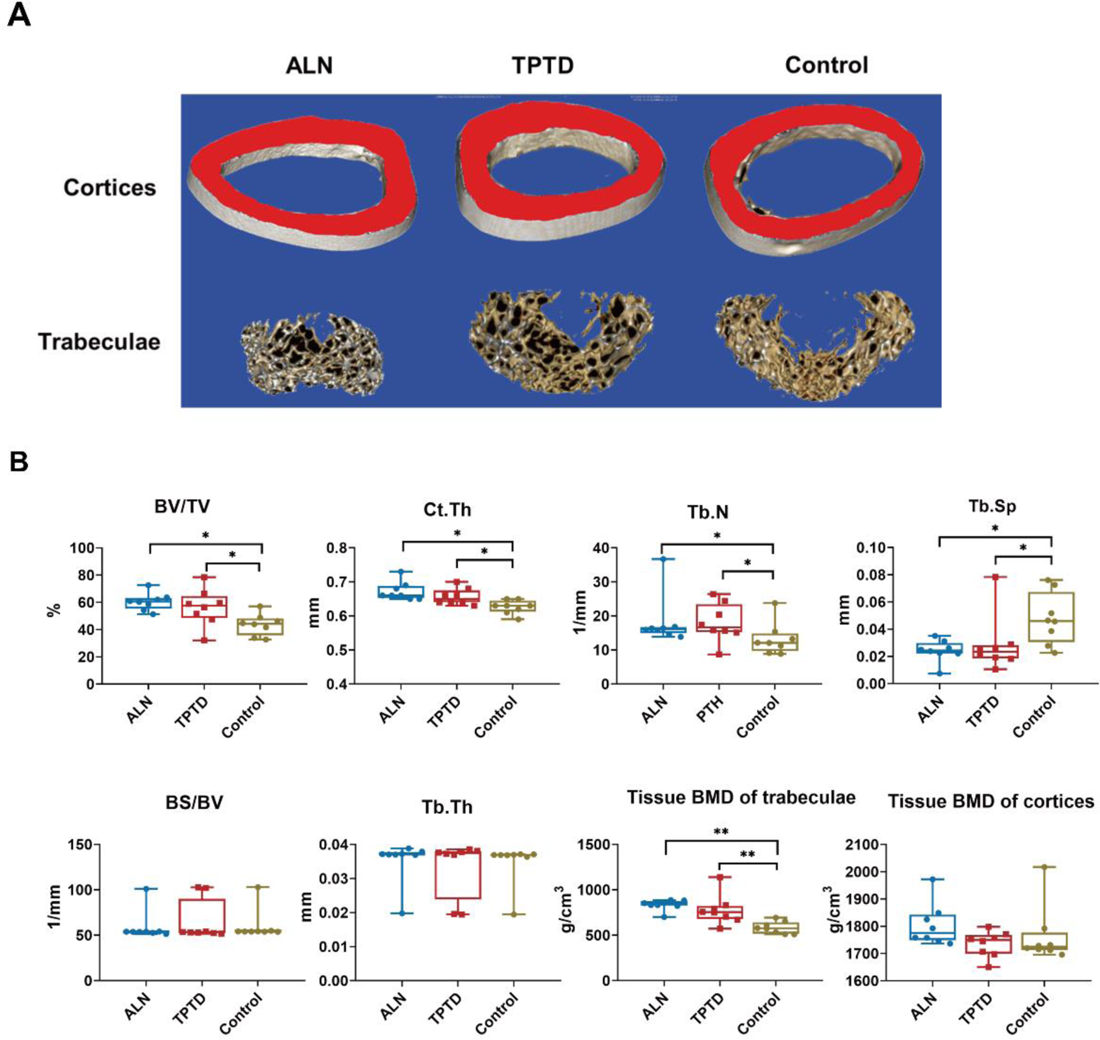
The efficacy of anti-osteoporotic treatment in *PLS3^E10-16del/0^* rats. Fig. 3A Three-dimensional reconstruction images of femurs after treatment Fig. 3B Microstructural parameters of femurs by micro-CT after treatment ALN: alendronate, TPTD: teriparatide, BV/TV: bone volume / tissue volume, BS/BV: bone surface area / bone volume, Tb.Th: trabecular thickness, Tb.N: trabecular number, Tb.Sp: trabecular separation, Ct.Th: cortical thickness, BMD: tissue bone mineral density Data are shown as the mean ± SD, evaluated by one-way ANOVA followed by Tukey’s post-hoc test. \**P* < 0.05; \*\**P* < 0.01; \*\*\**P* < 0.001.

**Fig. 4.**
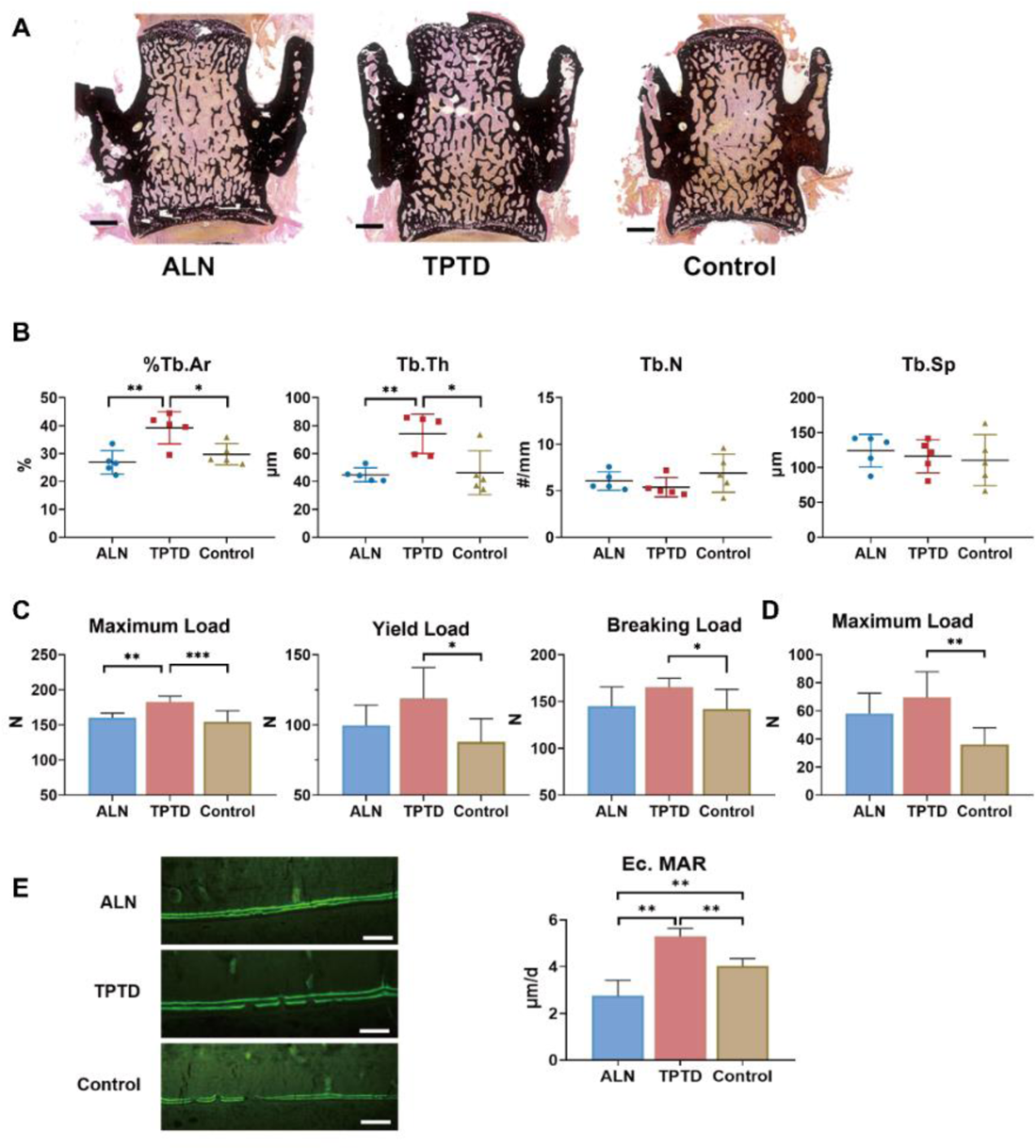
Changes in microarchitecture and strength after anti-osteoporotic treatment. Fig. 4A Typical images of unstained and uncalcified vertebra of *PLS3^E10-16del/0^*rats after treatment Scale bar = 1000 μm Fig. 4B Histomorphometric analysis of L4 after treatment %Tb.Ar: trabecular area, Tb.Th: trabecular thickness, Tb.N: trabecular number, Tb.Sp: trabecular separation Fig. 4C Effects of treatment on the mechanical strength of femoral diaphysis The diaphysis was subjected to three-point bending test to failure, which provided data on yield load, maximum load, breaking load. Fig. 4D Effects of treatment on the mechanical strength of L5 The vertebral body was subjected to indentation test to acquire maximum load. Fig. 4E Comparison of Ec.MAR in the tibial cortex among three treatment groups. (n > 3 per group) Scale bar = 100 μm ALN: alendronate, TPTD: teriparatide, control: saline. Ec.MAR: mineral apposition rate of endocortical surface of tibia cortex. Data are shown as the mean ± SD, evaluated by one-way ANOVA followed by Tukey’s post-hoc test. *P < 0.05; **P < 0.01; ***P < 0.001.

Moreover, TPTD treatment significantly increased maximum load (182.2 ± 8.7 N vs 154.2 ± 15.9 N, *P* < 0.001), yield load (117.5± 22.8 N vs 87.8 ± 16.6 N, *P* < 0.05) and breaking load (165.3± 9.5 N vs 142.2 ± 20.8 N, *P* < 0.01) of femur and maximum load of the fifth lumbar vertebrae (69.5± 18.3 N vs 32.1± 5.1 N, *P* < 0.01) than VEH group. (Fig.4C, 4D). The bone strength of femur and lumbar vertebrae was not significantly improved in ALN group.

## Discussion

Recent advancements in genetic research have uncovered that loss-of-function variants in *PLS3* can cause a monogenic X-linked early-onset osteoporosis and osteoporotic fractures, but the exact mechanism is unknown (Wolff et al., 2021). The optimal treatment regimen has not been established for *PLS3*-related osteoporosis. In the present study, we have generated a novel rat model with large fragment deletion in *PLS3*, and we systematically assessed bone microarchitecture, bone biomechanical property, BMD and bone remodeling for the first time in this novel rat model with patient-derived mutation. Interestingly, the newly generated *PLS3^E10-16del/0^* rat model displayed significantly impaired bone strength, decreased cortical thickness, and decreased mineral apposition rate of trabecular bone. We demonstrated that treatment with alendronate and teriparatide could significantly improve bone microstructure and increase BMD, and teriparatide could obviously improve the bone strength of *PLS3^E10-16del/0^* rat.

In this study, *PLS3^E10-16del/0^* rats displayed a bone-specific phenotype, including significantly impaired bone strength, decreased cortical bone thickness, and decreased mineral apposition rate, despite the ubiquitous presence of the mutation. Similar results were found in patients with various *PLS3* mutations and the *PLS3*-deficient mice model (Costantini et al., 2018; Fahiminiya et al., 2014; Hu et al., 2020; Kampe et al., 2017; Kannu et al., 2017; Laine et al., 2015; Lv et al., 2017; Neugebauer et al., 2018; van Dijk et al., 2013; Wang et al., 2020; Yorgan et al., 2020), which demonstrated that PLS3 had an indispensable role in bone metabolism. PLS3 is expressed in all solid tissues except hematopoietic cells and is involved in all processes dependent on F-actin dynamics. In bone, PLS3 was widely expressed in osteoblasts, osteoclasts, and osteocytes (Fahiminiya et al., 2014; Kamioka, Sugawara, Honjo, Yamashiro, & Takano-Yamamoto, 2004; Neugebauer et al., 2018). Bone histomorphometric analysis indicated that the quantities of bone cells were normal in *PLS3^E10-16del/0^* rats, which was consistent with previous reports (Neugebauer et al., 2018; Yorgan et al., 2020). However, dysfunction of bone cells in *PLS3* mutant mice and patients was found in the current and previous studies. Animal studies indicated PLS3 involvement in cytoskeletal actin bundling (Oprea et al., 2008), thus mediating mechanotransduction in osteocytes and further affecting cellular signal transduction between osteoblast and osteoclast, though the regulation was not confirmed in any studies (Pathak et al., 2020; van Dijk et al., 2013). Studies on patients’ bone biopsies collectively insinuated a role for PLS3 in bone matrix mineralization (Balasubramanian et al., 2018; Kampe et al., 2017; Kannu et al., 2017; Laine et al., 2015). Impaired osteoblastic bone mineralization induced by *PLS3* mutation was also observed in cell experiments and a murine model (Fahiminiya et al., 2014; Yorgan et al., 2020). F-actin-bundling ability or Ca^2+^ sensitivity would be disturbed after *PLS3* mutations (Schwebach et al., 2020), which would affect intracellular calcium concentrations that osteoblasts needed during differentiation and bone formation (Wang et al., 2018). In *PLS3^E10-16del/0^* rats, we also found an obvious reduction in MAR of trabecular bone, indicating PLS3 was possibly involved in regulating the mineralization of osteoblasts. More recently, experimental findings suggested the regulatory function of PLS3 in osteoclastogenesis and osteoclast function through influencing podosome organization (Neugebauer et al., 2018). Taken together, these observations indicated abnormal bone metabolism caused by *PLS3* was possibly attributed to dysfunctional F-actin dynamics. However, the exact roles of *PLS3* regulating cells of bone still needed to be further elucidated.

So far, the underlying molecular mechanism remained elusive. In *PLS3* mutation-positive patients, serum DKK1 concentrations were significantly elevated compared with WNT1 mutation-positive patients, which indicated impaired WNT signaling may involve in the occurrence of *PLS3*-related osteoporosis (Makitie et al., 2020). *PLS3-*deficient murine model exhibited decreased expression of *WNT16* (Yorgan et al., 2020)*. WNT16*-deficient mice displayed cortical bone defects with normal trabecular bone, with unchanged cortical MAR (Movérare-Skrtic et al., 2014). Both *PLS3* and *WNT16* could inhibit osteoclasts formation by inhibiting NF-κB activation and *Nfatc1* expression (Movérare-Skrtic et al., 2014; Neugebauer et al., 2018). Therefore, *PLS3* mutation could lead to increased osteoclasts differentiation through decreased *PLS3* and *WNT16* expression, which could lead to thin cortical bone. However, serum bone turnover biomarker levels of ALP and β-CTX were normal in our *PLS3^E10-16del/0^* rats and patients with *PLS3* mutations (Balasubramanian et al., 2018; Fahiminiya et al., 2014; Kampe et al., 2017; Laine et al., 2015). More in-depth studies were needed.

Since patients with *PLS3* mutation presented with low bone mineral density (BMD), peripheral fractures and multiple vertebral compression fractures, particularly in the thoracic spine (Hu et al., 2020; Lv et al., 2017; R. E. Mäkitie et al., 2020; van Dijk et al., 2013), it is of great clinical significance to establish an effective treatment regimen for *PLS3* related osteoporosis. Bisphosphonates such as alendronate are still recommended as the first-line antiresorptive therapy for osteoporosis. Teriparatide (TPTD), a recombinant human parathyroid hormone (1-34), is an extensively used bone anabolic drug for osteoporosis. It is noticed that the mechanisms of action are completely different between antiresorptive and osteoanabolic agents. Alendronate inhibits osteoclasts and bone resorption mainly via the mevalonate pathway (Cremers, Drake, Ebetino, Bilezikian, & Russell, 2019). On the other hand, osteoanabolic TPTD stimulates bone formation by affecting protein kinases, MAP-kinase, phospholipases, as well as the WNT-signaling pathway (Canalis, Giustina, & Bilezikian, 2007). In this study, alendronate and TPTD were all effective in increasing BMD and improving bone microstructure of *PLS3^E10-16del/0^* rats. Our results were consistent with the efficacy of ALN and TPTD in patients with PLS3-related EOOP (Fratzl-Zelman et al., 2021; Hu et al., 2020; Lv et al., 2017; Valimaki et al., 2017; van Dijk et al., 2013). Interestingly, TPTD treatment obviously improved the bone mechanical strength of femora and vertebrae of *PLS3^E10-16del/0^* rats, consistent with the improvement of bone microstructure of the above sites. However, we did not observe that alendronate significantly improved bone biomechanical properties of *PLS3^E10-16del/0^*rats, which may be related to the small dose of ALN and the short treatment time. In another study, treatment with ALN at a dose of 30 ug/kg/d for 12 weeks significantly improved stiffness of a murine model of osteogenesis imperfecta (McCarthy et al., 2002). It is necessary to carry out studies with a larger dose and longer time treatment of ALN to clarify its effects on bone strength of *PLS3^E10-16del/0^* rats.

Although the precise molecular mechanisms of *PLS3* mutation inducing early-onset osteoporosis needed to be clarified, we confirmed that *PLS3* played a critical role in regulating bone metabolism and maintaining the integrity of bone structure and mechanical properties in a new generated transgenic rat model with a large fragment deletion of *PLS3*. Our results demonstrated that teriparatide and alendronate are effective drugs for the first time in a transgenic rat model of *PLS3*-related osteoporosis, which provides valuable experimental evidence for the treatment of *PLS3*-related EOOP. However, there were a few limitations. Since *PLS3* was widely expressed in the body, we did not conduct morphometrical and functional analyses of other tissues and organs. We did not conduct in-depth research on the signal pathways regulating bone metabolism, such as WNT/β-catenin, OPG-RANK-RANKL pathway, and so on, which were helpful to clarify the pathogenesis of this EOOP.

In conclusion, *PLS3* plays a major role in bone metabolism and bone integrity. Impaired bone microstructure and bone strength were prominent characteristics of rats carrying hemizygous E10-16del mutation of *PLS3*, of which the exact molecular pathogenesis still needs further study. Alendronate and teriparatide treatment can increase BMD and improve the bone microstructure of rats with early-onset osteoporosis induced by *PLS3* mutation.

## Material and methods

### Generation of *PLS3* knock-out rat model and genetic identification

General *PLS3* knock-out rat model was independently generated using the CRISPR/Cas9 system at the Institute of Laboratory Animal Sciences, Chinese Academy of Medical Sciences & Peking Union Medical College. Five specific guide RNAs were used to target the exon 10-16 of *PLS3* (Table 1), which were annealed and cloned into the PUC57-gRNA expression vector (Addgene 51132, Cambridge, MA, USA) with a T7 promoter. In vitro, transcription of gRNA template was accomplished using the MEGAshortscript Kit (AM1354, Ambion). The pST1374-NLS-flag-linker-Cas9 vector (Addgene 44758) was linearized using the Age I enzyme and transcribed with a T7 Ultra Kit (Ambion, AM1345). After being purified with the MEGAclear Kit (AM1908, Ambion), a mixture of transcribed Cas9 mRNA and gRNA was microinjected into the cytoplasm of zygotes of Sprague Dawley (SD) rats which were obtained commercially from Beijing Vital River Laboratory Animal Technology Co., Ltd.

**Table 1.**
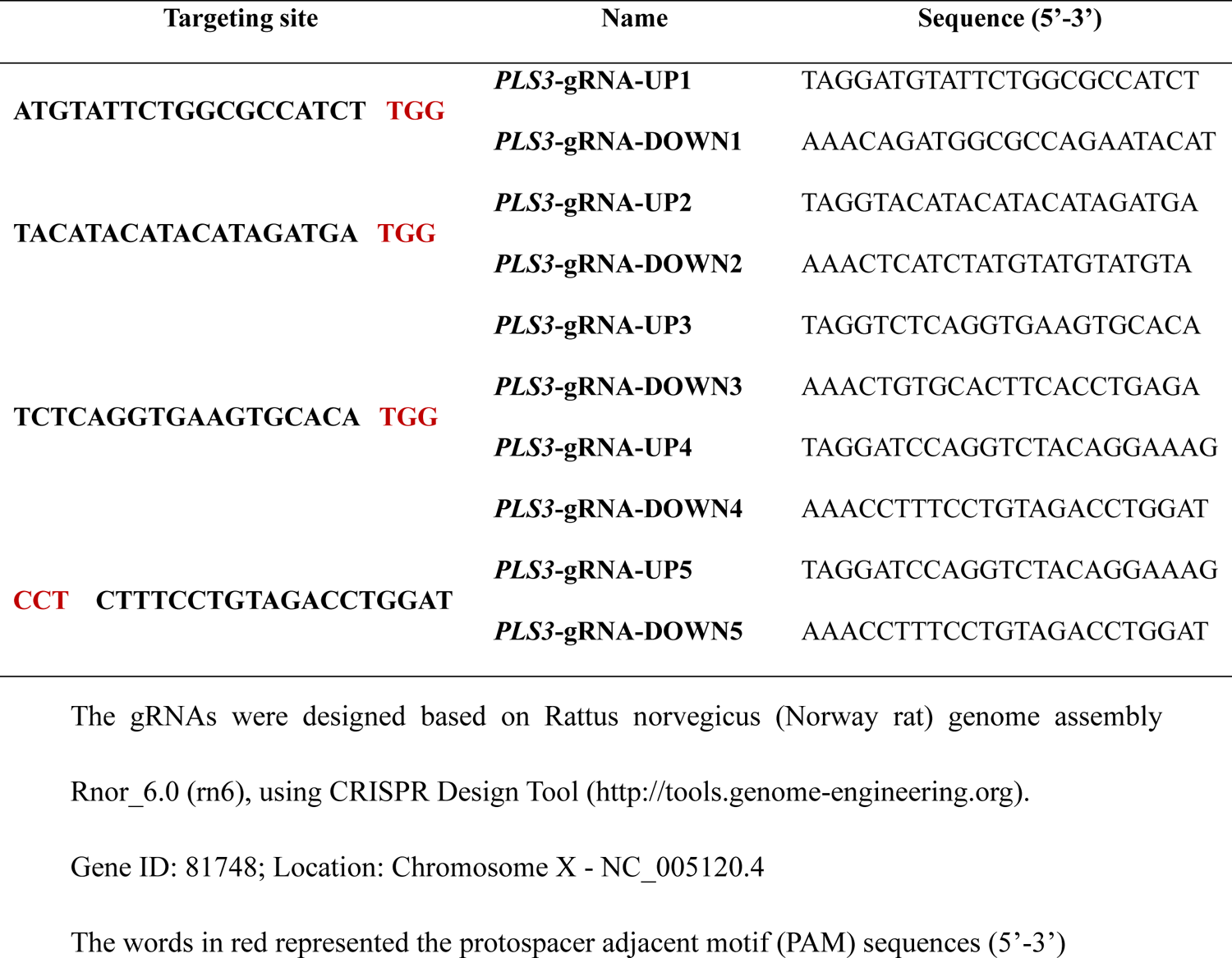
Small guide RNA (gRNA) sequence for *PLS3* gene knockout

A male founder was backcrossed with wild-type rats to generate female heterozygous knock-out rats (*PLS3^E10-16del^*^/+^), which were subsequently crossed with wild-type male. Being located on the X chromosome, PLS3-induced osteoporosis was more severe in males than females, male hemizygous knock-out rats (*PLS3 ^E10-16del^*^/0^) of the offspring were selected to be further studied. Since osteoporosis associated with *PLS3* mutations had its disease onset early in childhood and adolescence, and rats reached the peak bone mass at 6-9 months of age, we chose rats at the age of 3, 6, and 9 months for further investigation. All rats were registered and preserved on a SD genetic background in a specific pathogen-free (SPF) environment without surpassing 6 animals in ventilated cages. Rats were provided with standard chow and water ad libitum.

Genomic DNA was isolated from tail snips using E.Z.N.A.®Tissue DNA Kit (Omega Bio-tek, Norcross, GA, US). Genotyping was performed using polymerase chain reaction (PCR) amplification and Sanger sequencing. The allele-specific primers were as follows: forward 5’-CTTCATTCCCTTTGCACGTT-3’ (84092-84111 nt) and reverse 5’-TTTACTTACATGAAGACCCCT-3’ (94242-94262 nt) for *PLS3* knock-out (KO) allele, and a 652bp PCR production would be generated. Forward 5’-CCCATAAGTTGTTCCTTGATTTCC-3’ (88350-88373 nt) and reverse 5’-CACTGCCTGAATAAGACCCACTC-3’ (88777-88799 nt) for wild-type allele, a 450bp PCR production would be generated. Thermal cycling conditions consisted of an initial denaturation at 95 ℃ for 3 min, followed by 38 cycles at 95℃ for 30s, 52-58℃ for 30s, and 72℃ for 1 min. The PCR products were sequenced directly.

### Treatment

A total of 24 *PLS3^E10-16del^*^/0^ male rats with three months old were randomized either to vehicle (VEH), alendronate (ALN) or teriparatide (TPTD) therapy (n=8 in each group) for 8 weeks. Alendronate 1.0 ug/kg body weight was subcutaneously injected into rats in ALN group daily (alendronate sodium pure dry, provided by Merck and Co., Inc., Rahway, NJ, USA) (Diab, Wang, Reinwald, Guldberg, & Burr, 2011; Iwata, Li, Follet, Phipps, & Burr, 2006). In the TPTD group, the agent (Recombinant Human Parathyroid Hormone 1-34, provided by salubris biotherapeutics, Inc., Shenzhen, China) was administered s.c. at the dose of 40ug/kg body weight, five times weekly (Komrakova et al., 2010; Lane et al., 1996). Rats in the VEH group received vehicle injections (0.9% saline dosed at the same volume of 1 mL/kg body weight) s.c. daily, and thus acted as control groups.

All animal experiments were approved by the Institutional Animal Care and Use Committee of the Peking Union Medical College Hospital (XHDW-2021-027).

### Bone microstructure assessment

The left femur was fixed in 4% paraformaldehyde (PFA), of which microarchitecture was assessed by micro-computed tomography (μCT) (Inveon MM CT, Siemens, Erlangen, Germany) according to the recommended protocol (Bouxsein et al., 2010). In vitro scans were operated with an X-ray tube voltage of 60 kV, a current of 400 μA, an exposure time of 800 ms and a voxel size of 20 μm. The region of interest (ROI) for trabecular bone was drawn in the distal epiphysis, starting 1.5 mm below the growth plate and extending 100 slices to proximal end. Cortical bone was analyzed in a 1000-μm-long volume situated in the middle of the diaphysis. BMD, bone volume / tissue volume (BV/TV), bone surface area / bone volume (BS/BV), cortical thickness (Ct.Th), trabecular thickness (Tb.Th), trabecular number (Tb.N) and trabecular separation (Tb.Sp) were measured. The Inveon Research Workplace software (Siemens) was used for reconstruction and analysis of two-dimensional (2D) and 3D image.

### Assessment of biomechanical properties of bone

The right femur and the fifth lumbar vertebrae (L_5_) were wrapped in saline-soaked gauze and stored at –20 °C, which were thawed at room temperature 2 hours before the mechanical test. Three-point binding tests (TPBT) and indentation testing were implemented on a fatigue-testing machine (BOSE ElectroForce 3200, TA Instruments, New Castle, DE, USA). The right femurs were placed in a customized holder with the span between supports fixed at 2 cm, and the crosshead lowered at a constant speed of 1 mm/min until the femur fractured. From the load-displacement curves, stiffness, yield load, maximum load, breaking load, post-yield displacement and work-to-fracture were generated.

Force-displacement measurement was performed on the fifth lumbar vertebral bodies. The probe was custom-made (d = 1.0mm) and fixed on the holder with a sensor. Vertebral arch and end-plates were removed to obtain a specimen with planoparallel ends. Each specimen was placed in embedding plastic and oriented to support the test surface perpendicular to the indenter. The resulting force-displacement curve was shown together, and maximum load were measured.

### Measurement of bone metabolic markers

Under abdominal anesthesia, blood samples were collected via cardiac puncture. Concentrations of serum calcium (Ca), phosphorus (P), alkaline phosphatase (ALP, a bone formation marker) were measured by an automated chemistry analyzer (AU5800, Beckman Coulter Inc., Brea CA, USA). Serum level of C-telopeptide of type Ⅰ collagen (β-CTX, bone resorption marker) was measured by ELISA (Cat# CSB-E12776r, Cusabio Biotech Co., Wuhan, China).

### Analysis of histology and histomorphometry

All rats received intraperitoneal injection of calcein (10mg/kg body mass, Sigma-Aldrich, Co., St. Louis, MO, USA) for histomorphometric analysis on the second and the 6th day before euthanization. The left femur was decalcified after μCT scanning and were embedded in paraffin and cut into 4-μm sections using a microtome (Leica RM2016, Leica Microsystems). Hematoxylin & eosin (H&E) staining and tartrate-resistant acid phosphatase (TRAP) staining (Servicebio, Cat# G1050) were performed. The osteoclast number per bone perimeter (N.Oc/B.Pm), osteocyte number per bone area (N.Ot/B.Ar), and osteoblast number/bone perimeter (N.Ob/B.Pm) were calculated.

The right tibia and the fourth vertebrae (L_4_) were fixed in 70% alcohol and embedded in modified methyl methacrylate without decalcification. The embedded samples were cut into 10-μm thick sections, which were de-plasticized and stained with von-kossa stain kit (Servicebio, Cat# G1043) to calculate trabecular area (%Tb.Ar), Tb.Th, Tb.N, and Tb.Sp. For unstained slices, the mineral apposition rate (MAR) was calculated by dividing the distance between the two calcein labels by the inter-labeling period. Analysis was performed with Image J software according to the recommendation of ASBMR (Dempster et al., 2013).

## Statistical Analysis

All experiments were repeated at least thrice independently. Results were reported as means ± SD (standard difference). Comparisons of parameters between transgenic and wild type rats were completed using Student’s t test. Parameters of rats at different ages or in different treatment groups were compared with one-way ANOVA followed by Tukey’s post-hoc test. Statistical analysis was performed using SPSS Statistics 26.0 (IBM, Armonk, NY, USA), GraphPad Prism 8 (Statcon). Statistical significance was determined when *P* values equal to or less than 0.05.

## Funding Information

This work is supported by National Key R&D Program of China (2018YFA0800801, 2021YFC2501704), CAMS Innovation Fund for Medical Sciences (CIFMS)(2021-I2M-C&T-B-007, 2021-I2M-1-051), National Natural Science Foundation of China (No.81873668, 82070908), and Beijing Natural Science Foundation (7202153)

## Declaration of interest

No competing interests declared.

## Author contributions

Conceptualization, Investigation, Visualization, Writing - original draft. Conceptualization, Investigation, Visualization.

Conceptualization, Investigation. Investigation, Methodology.

Resources, Investigation, Methodology. Conceptualization, Investigation.

Conceptualization, Investigation. Resources, Supervision.

Resources, Supervision.

Resources, Writing - review & editing. Writing - review & editing.

Conceptualization, Writing - original draft, Supervision, Funding acquisition.

## Ethics

This study was performed in strict accordance with the recommendations in the Guide for the Care and Use of Laboratory Animals of the National Institutes of Health. All animal experiments were approved by the Institutional Animal Care and Use Committee of the Peking Union Medical College Hospital (XHDW-2021-027). Every effort was made to minimize pain and suffering by providing support when necessary and choosing ethical endpoints.

## Data availability

All data generated or analyzed during this study are included in the manuscript and supporting source data files.

## Supplemental information

Supplemental data include three figures.

---

**Supplementary Fig. 1A.**
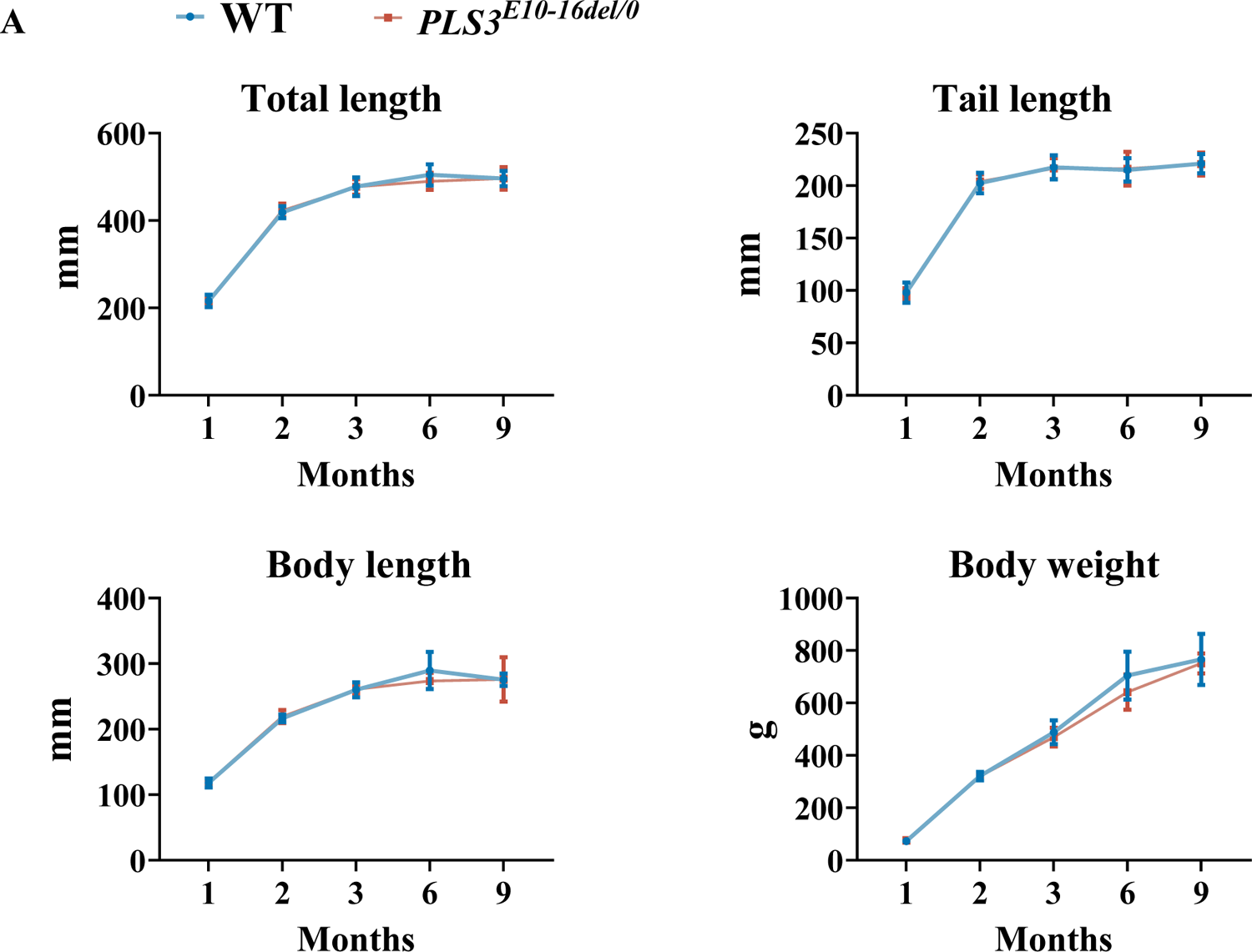
Growth curve of *PLS3^E10-16del/0^* and WT rats. WT: wild-type

**Supplementary Fig. 2A.**
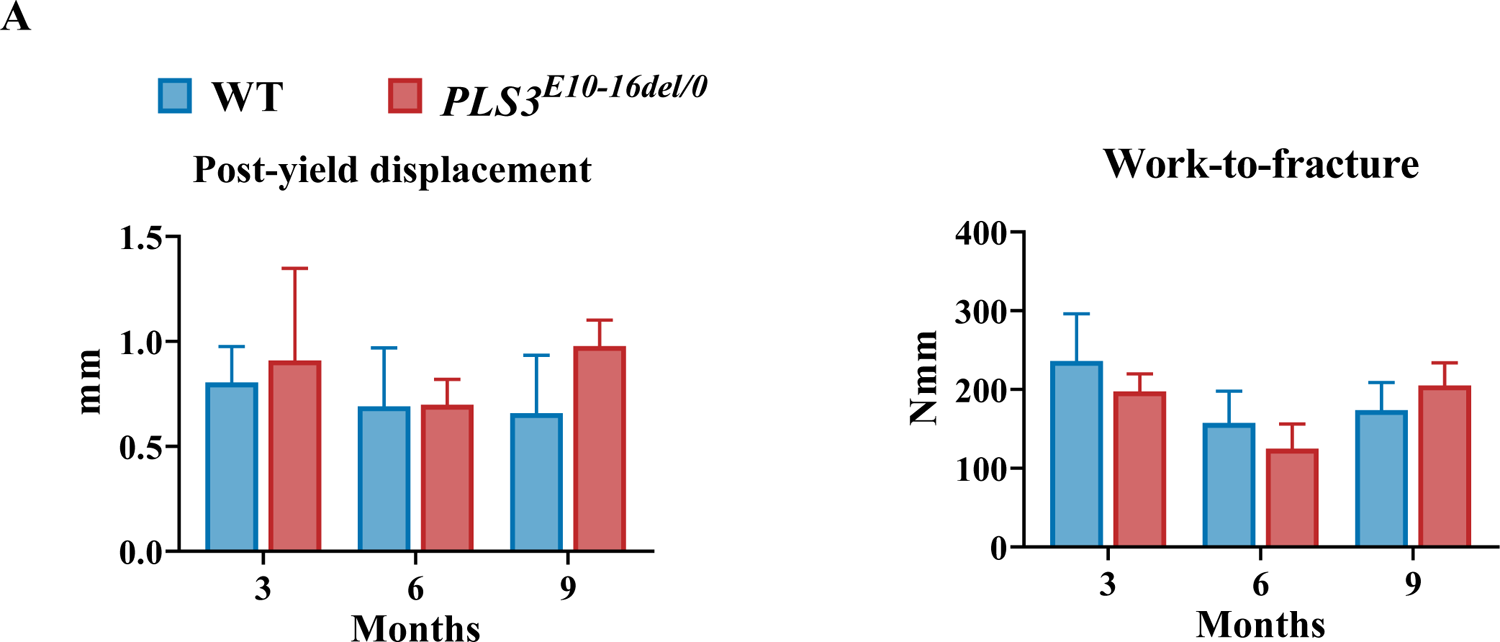
Mechanical three-point bending tests of femora from *PLS3^E10-16del/0^* and WT rats (n= 5-8 per group) WT: wild-type

**Supplementary Fig. 2B.**
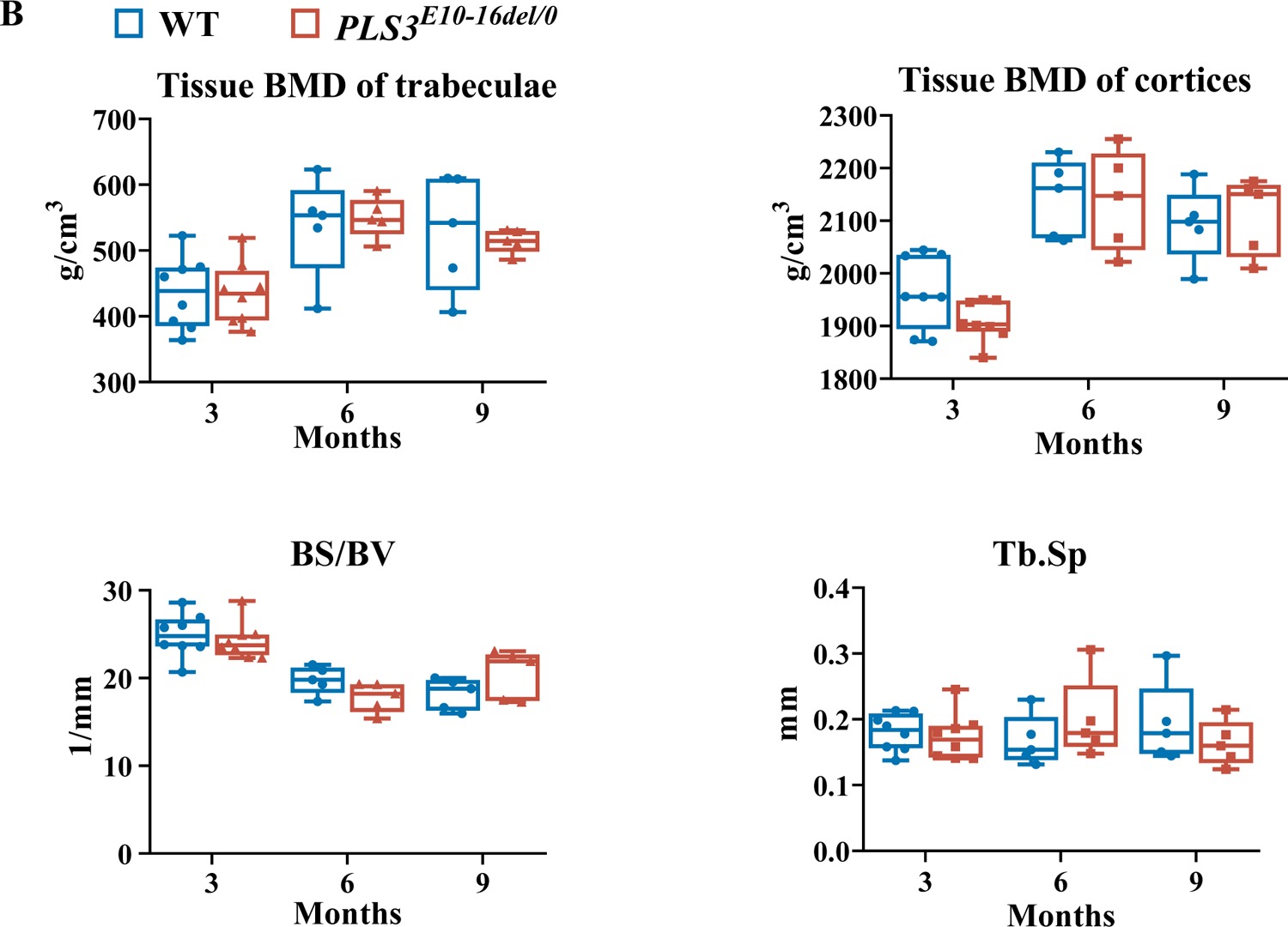
Micro-CT assessment of the distal femur from *PLS3^E10-16del/0^* and WT rats BS/BV: bone surface/bone volume, Tb.Sp: trabecular separation WT: wild-type

**Supplementary Fig2A.**
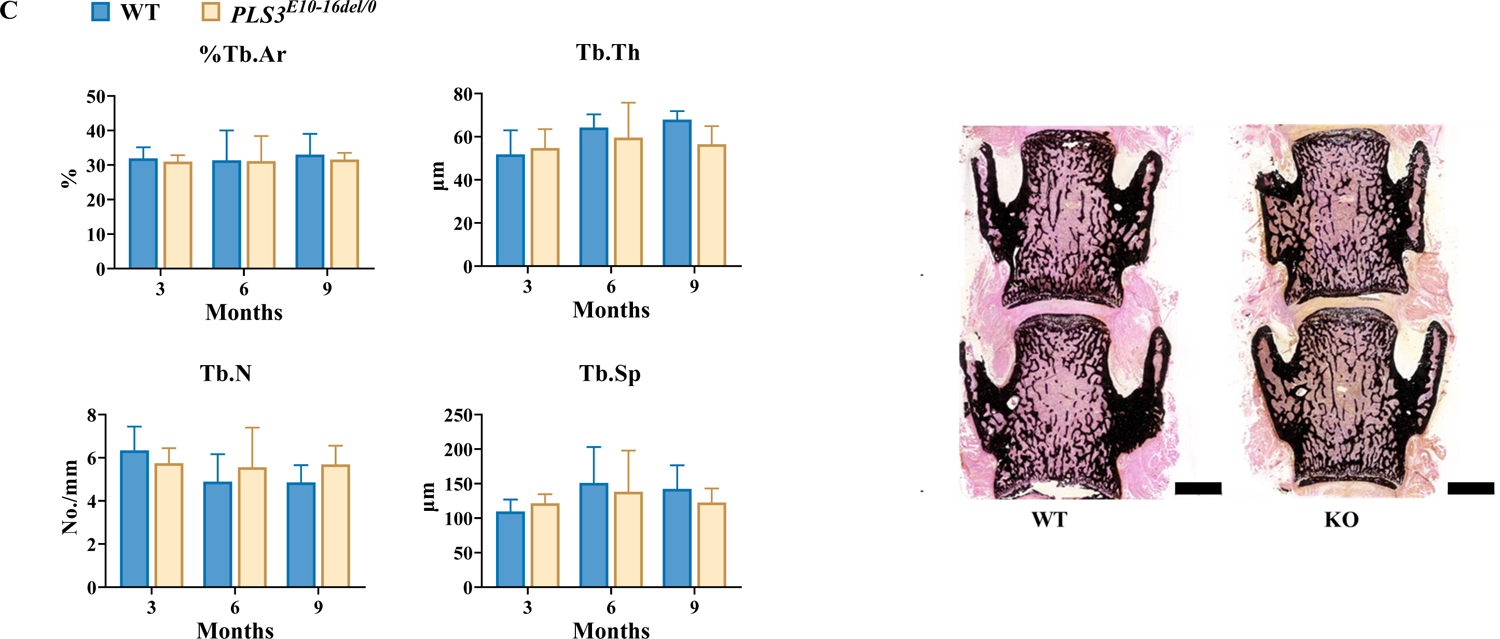
Histomorphometric evaluation of L4 from *PLS3^E10-16del/0^* and WT rats (n= 5-7 per group) %Tb.Ar: trabecular area, Tb.Th: trabecular thickness, Tb.N: trabecular number, Tb.Sp: trabecular separation Scale bar = 2000 μm WT: wild-type

**Supplementary Fig2D.**
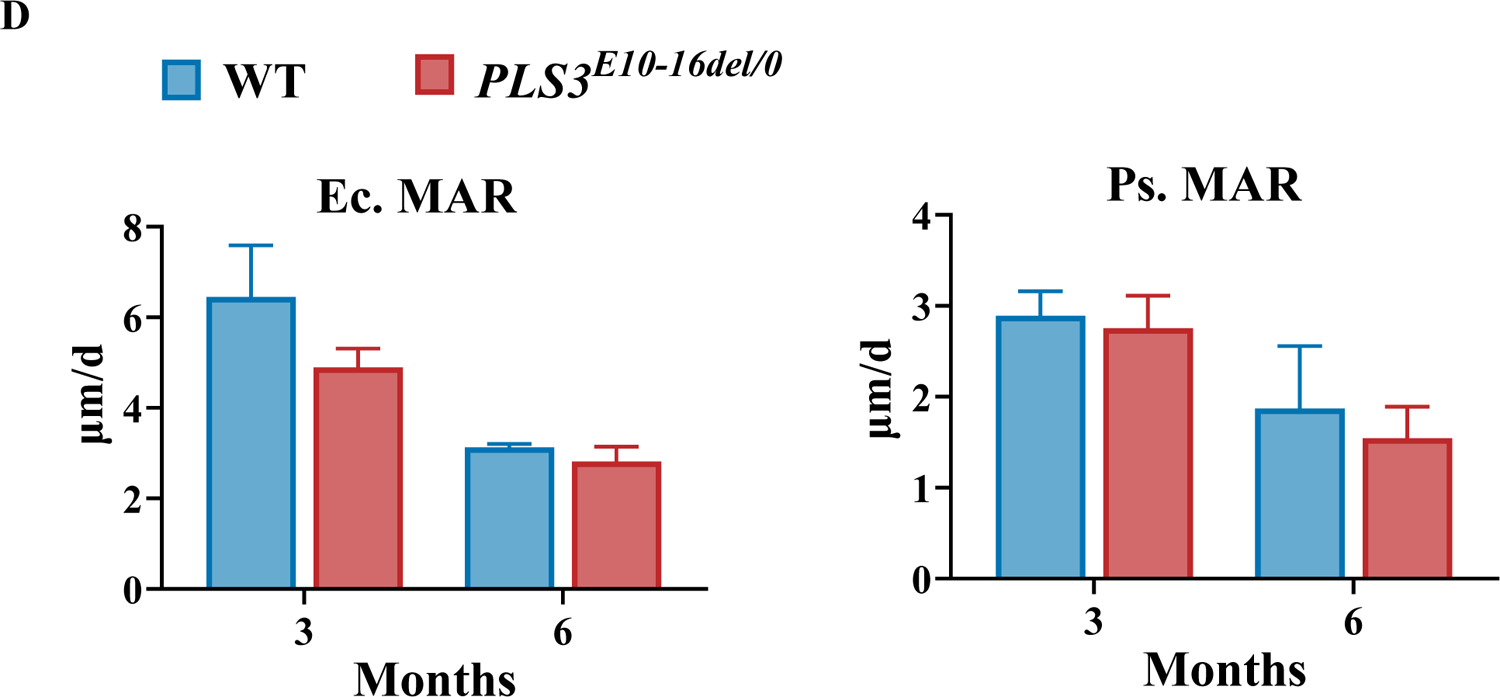
Analysis of MAR in tibial cortices of *PLS3^E10-16del/0^* rats. Ec.MAR: mineral apposition rate of endocortical surface of tibia cortex Ps.MAR: mineral apposition rate of periosteal surface of tibia cortex WT: wild-type,

**Supplementary Fig2E.**
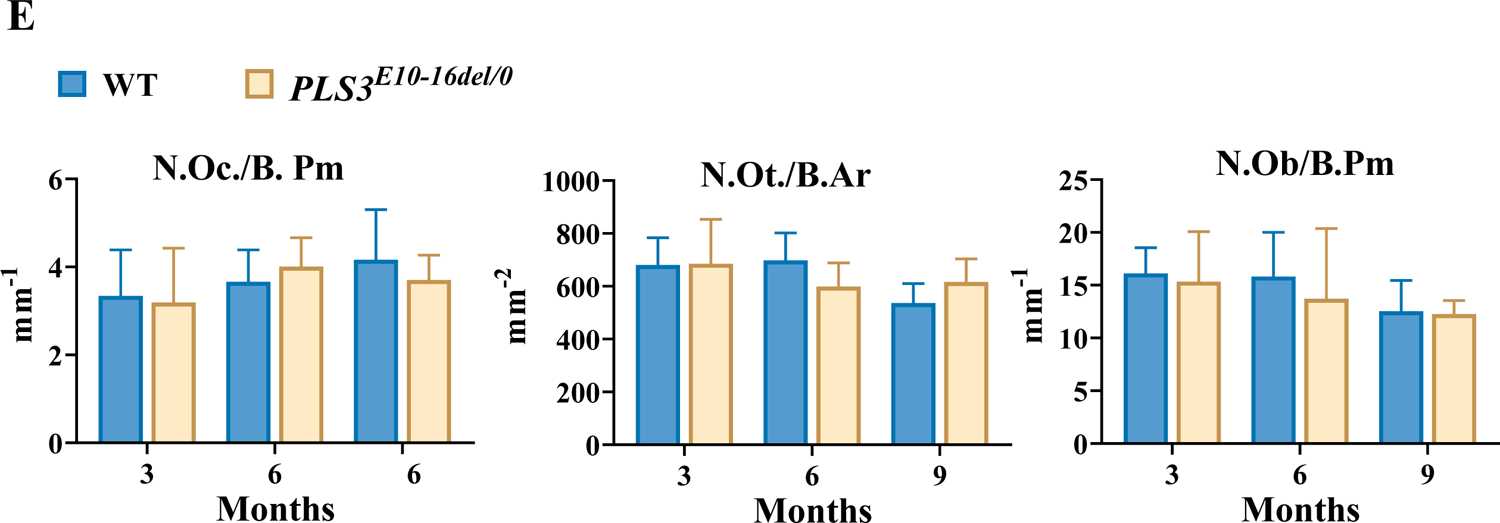
Measurements of osteoclasts, osteocytes and osteoblasts of *PLS3^E10-16del/0^* rats (n= 5-8 per group) N.Oc./B.Pm: osteoclast number per bone perimeter, N.Ot./B.Ar:osteocyte number per bone area; osteoblast number/bone perimeter

**Supplementary Fig. 2F.**
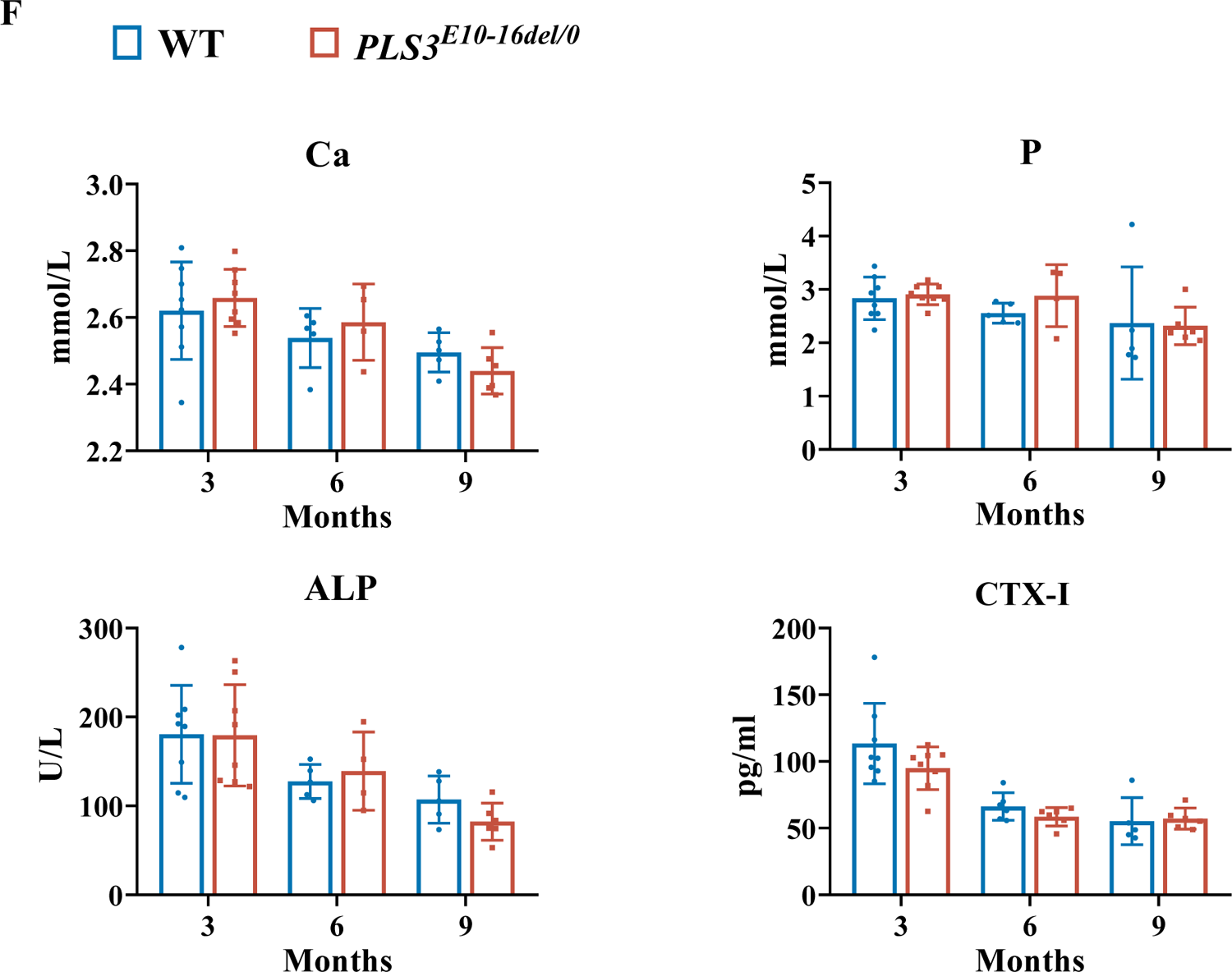
Serum levels of bone metabolic markers in *PLS3^E10-16del/0^* and WT rats. Ca: calcium, P: phosphorus, β-CTX: β-C-telopeptide of type Ⅰ collagen, bone resorption marker, ALP: alkaline phosphatase. ELISAs were used to determine the levels of CTX-I. An automated chemistry analyzer was used to measure the levels of the Ca, P and ALP.

**Supplementary Fig. 3A.**
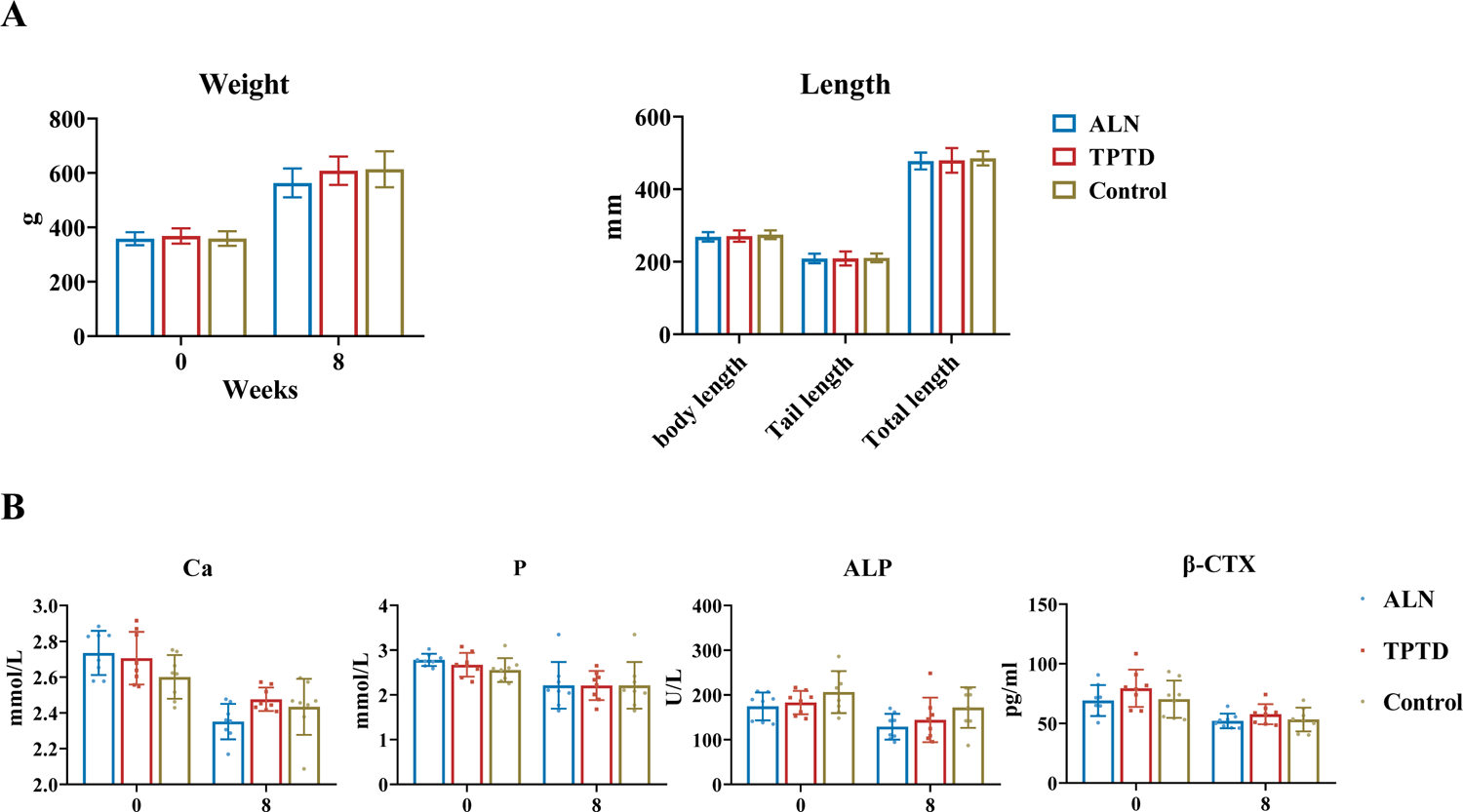
Body mass and length of rats at the initiation and end of the treatment (n=8 per group) Body mass were comparable among three groups at the initiation and end of the treatment. No differences were found in body length, tail length and total length of rats at the completion of 2 months of therapy. ALN: alendronate group; TPTD: teriparatide **Supplementary Fig. 3B** Serum levels of bone metabolic markers during treatment Ca: calcium, P: phosphorus, β-CTX: β-C-telopeptide of type Ⅰ collagen, bone resorption marker, ALP: alkaline phosphatase. Bone metabolic markers were measured at 0 and 8weeks of treatment

**Supplementary Fig. 3C.**
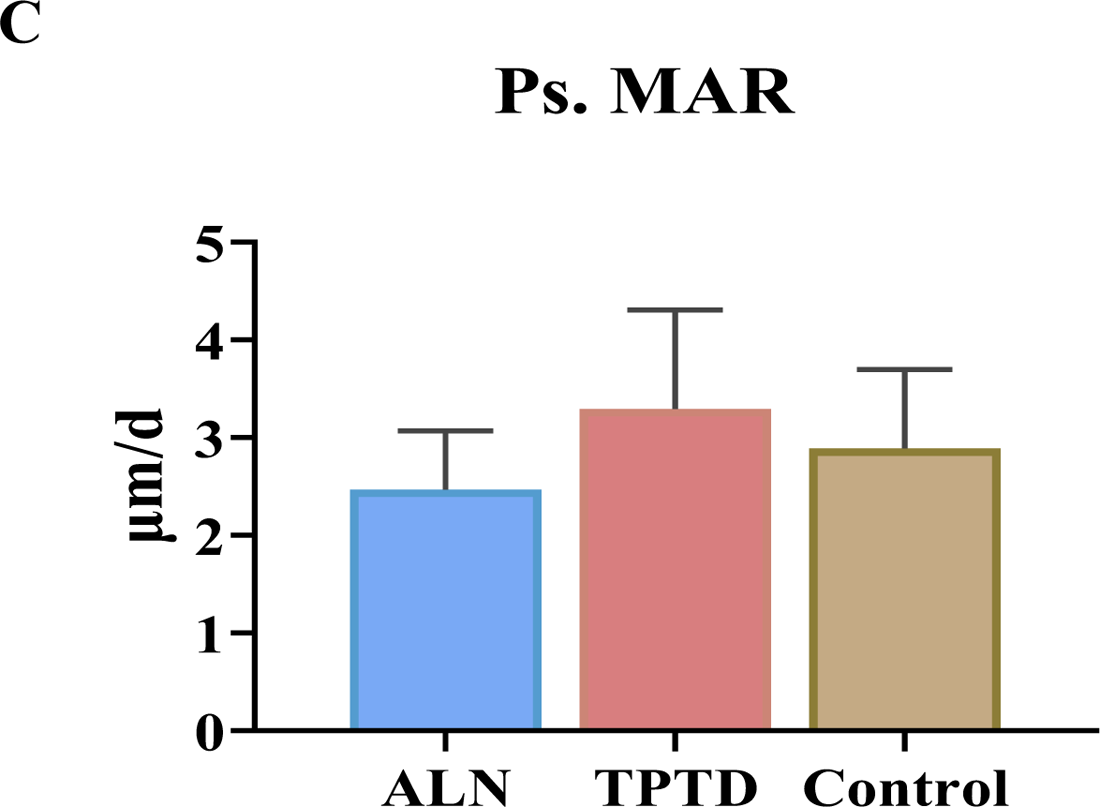
Comparison of Ps.MAR in the tibial cortex among three treatment groups. (n > 3 per group) Ps.MAR: mineral apposition rate of periosteal surface of tibia cortex.

## Notes

### Competing Interest Statement

The authors have declared no competing interest.

## Reference

1. Balasubramanian, M., Fratzl-Zelman, N., O’Sullivan, R., Bull, M., Fa Peel, N., Pollitt, R. C., … Bishop, N. J. (2018). Novel PLS3 variants in X-linked osteoporosis: Exploring bone material properties. Am J Med Genet A, 176(7), 1578–1586. doi:10.1002/ajmg.a.38830

2. Bouxsein, M. L., Boyd, S. K., Christiansen, B. A., Guldberg, R. E., Jepsen, K. J., & Müller, R. (2010). Guidelines for assessment of bone microstructure in rodents using micro-computed tomography. Journal of Bone and Mineral Research, 25(7), 1468–1486. doi:10.1002/jbmr.141

3. Brlek, P., Antičević, D., Molnar, V., Matišić, V., Robinson, K., Aradhya, S., … Primorac, D. (2021). X-Linked Osteogenesis Imperfecta Possibly Caused by a Novel Variant in PLS3. Genes (Basel), 12(12), 1851. doi:10.3390/genes12121851

4. Canalis, E., Giustina, A., & Bilezikian, J. P. (2007). Mechanisms of anabolic therapies for osteoporosis. N Engl J Med, 357(9), 905–916. doi:10.1056/NEJMra067395

5. Cohen, A., Hostyk, J., Baugh, E. H., Buchovecky, C. M., Aggarwal, V. S., Recker, R. R., … Shane, E. (2022). Whole exome sequencing reveals potentially pathogenic variants in a small subset of premenopausal women with idiopathic osteoporosis. Bone, 154, 116253. doi:10.1016/j.bone.2021.116253

6. Costantini, A., Krallis, P., Kampe, A., Karavitakis, E. M., Taylan, F., Makitie, O., & Doulgeraki, A. (2018). A novel frameshift deletion in PLS3 causing severe primary osteoporosis. J Hum Genet, 63(8), 923–926. doi:10.1038/s10038-018-0472-5

7. Cremers, S., Drake, M. T., Ebetino, F. H., Bilezikian, J. P., & Russell, R. G. G. (2019). Pharmacology of bisphosphonates. Br J Clin Pharmacol, 85(6), 1052–1062. doi:10.1111/bcp.13867

8. Dempster, D. W., Compston, J. E., Drezner, M. K., Glorieux, F. H., Kanis, J. A., Malluche, H., … Parfitt, A. M. (2013). Standardized nomenclature, symbols, and units for bone histomorphometry: a 2012 update of the report of the ASBMR Histomorphometry Nomenclature Committee. Journal of Bone and Mineral Research, 28(1), 2–17. doi:10.1002/jbmr.1805

9. Diab, T., Wang, J., Reinwald, S., Guldberg, R. E., & Burr, D. B. (2011). Effects of the combination treatment of raloxifene and alendronate on the biomechanical properties of vertebral bone. Journal of Bone and Mineral Research, 26(2), 270–276. doi:10.1002/jbmr.197

10. Fahiminiya, S., Majewski, J., Al-Jallad, H., Moffatt, P., Mort, J., Glorieux, F. H., … Rauch, F. (2014). Osteoporosis caused by mutations in PLS3: clinical and bone tissue characteristics. Journal of Bone and Mineral Research, 29(8), 1805–1814. doi:10.1002/jbmr.2208

11. Fratzl-Zelman, N., Wesseling-Perry, K., Makitie, R. E., Blouin, S., Hartmann, M. A., Zwerina, J., … Makitie, O. (2021). Bone material properties and response to teriparatide in osteoporosis due to WNT1 and PLS3 mutations. Bone, 146, 115900. doi:10.1016/j.bone.2021.115900

12. Hu, J., Li, L. J., Zheng, W. B., Zhao, D. C., Wang, O., Jiang, Y., … Xia, W. (2020). A novel mutation in PLS3 causes extremely rare X-linked osteogenesis imperfecta. Mol Genet Genomic Med, 8(12), e1525. doi:10.1002/mgg3.1525

13. Iwata, K., Li, J., Follet, H., Phipps, R. J., & Burr, D. B. (2006). Bisphosphonates suppress periosteal osteoblast activity independently of resorption in rat femur and tibia. Bone, 39(5), 1053–1058. doi:10.1016/j.bone.2006.05.006

14. Kamioka, H., Sugawara, Y., Honjo, T., Yamashiro, T., & Takano-Yamamoto, T. (2004). Terminal differentiation of osteoblasts to osteocytes is accompanied by dramatic changes in the distribution of actin-binding proteins. Journal of Bone and Mineral Research, 19(3), 471–478. doi:10.1359/jbmr.040128

15. Kampe, A. J., Costantini, A., Levy-Shraga, Y., Zeitlin, L., Roschger, P., Taylan, F., … Makitie, O. (2017). PLS3 Deletions Lead to Severe Spinal Osteoporosis and Disturbed Bone Matrix Mineralization. Journal of Bone and Mineral Research, 32(12), 2394–2404. doi:10.1002/jbmr.3233

16. Kannu, P., Mahjoub, A., Babul-Hirji, R., Carter, M. T., & Harrington, J. (2017). PLS3 Mutations in X-Linked Osteoporosis: Clinical and Bone Characteristics of Two Novel Mutations. Horm Res Paediatr, 88(3-4), 298–304. doi:10.1159/000477242

17. Komrakova, M., Stuermer, E. K., Werner, C., Wicke, M., Kolios, L., Sehmisch, S., … Stuermer, K. M. (2010). Effect of human parathyroid hormone hPTH (1-34) applied at different regimes on fracture healing and muscle in ovariectomized and healthy rats. Bone, 47(3), 480–492. doi:10.1016/j.bone.2010.05.013

18. Laine, C. M., Wessman, M., Toiviainen-Salo, S., Kaunisto, M. A., Mayranpaa, M. K., Laine, T., … Makitie, O. (2015). A novel splice mutation in PLS3 causes X-linked early onset low-turnover osteoporosis. Journal of Bone and Mineral Research, 30(3), 510–518. doi:10.1002/jbmr.2355

19. Lane, N. E., Kimmel, D. B., Nilsson, M. H., Cohen, F. E., Newton, S., Nissenson, R. A., & Strewler, G. J. (1996). Bone-selective analogs of human PTH(1-34) increase bone formation in an ovariectomized rat model. Journal of Bone and Mineral Research, 11(5), 614–625. doi:10.1002/jbmr.5650110509

20. Lv, F., Ma, M., Liu, W., Xu, X., Song, Y., Li, L., … Li, M. (2017). A novel large fragment deletion in PLS3 causes rare X-linked early-onset osteoporosis and response to zoledronic acid. Osteoporos Int, 28(9), 2691–2700. doi:10.1007/s00198-017-4094-0

21. Mäkitie, O., & Zillikens, M. C. (2022). Early-Onset Osteoporosis. Calcif Tissue Int, 110(5), 546–561. doi:10.1007/s00223-021-00885-6

22. Makitie, R. E., Kampe, A., Costantini, A., Alm, J. J., Magnusson, P., & Makitie, O. (2020). Biomarkers in WNT1 and PLS3 Osteoporosis: Altered Concentrations of DKK1 and FGF23. Journal of Bone and Mineral Research. doi:10.1002/jbmr.3959

23. Mäkitie, R. E., Niinimäki, T., Suo-Palosaari, M., Kämpe, A., Costantini, A., Toiviainen-Salo, S., … Mäkitie, O. (2020). PLS3 Mutations Cause Severe Age and Sex-Related Spinal Pathology. Front Endocrinol (Lausanne), 11, 393. doi:10.3389/fendo.2020.00393

24. McCarthy, E. A., Raggio, C. L., Hossack, M. D., Miller, E. A., Jain, S., Boskey, A. L., & Camacho, N. P. (2002). Alendronate treatment for infants with osteogenesis imperfecta: demonstration of efficacy in a mouse model. Pediatric Research, 52(5), 660–670. doi:10.1203/00006450-200211000-00010

25. Movérare-Skrtic, S., Henning, P., Liu, X., Nagano, K., Saito, H., Börjesson, A. E., … Ohlsson, C. (2014). Osteoblast-derived WNT16 represses osteoclastogenesis and prevents cortical bone fragility fractures. Nat Med, 20(11), 1279–1288. doi:10.1038/nm.3654

26. Neugebauer, J., Heilig, J., Hosseinibarkooie, S., Ross, B. C., Mendoza-Ferreira, N., Nolte, F., … Wirth, B. (2018). Plastin 3 influences bone homeostasis through regulation of osteoclast activity. Hum Mol Genet, 27(24), 4249–4262. doi:10.1093/hmg/ddy318

27. Oprea, G. E., Kröber, S., McWhorter, M. L., Rossoll, W., Müller, S., Krawczak, M., … Wirth, B. (2008). Plastin 3 is a protective modifier of autosomal recessive spinal muscular atrophy. Science, 320(5875), 524–527. doi:10.1126/science.1155085

28. Pathak, J. L., Bravenboer, N., & Klein-Nulend, J. (2020). The Osteocyte as the New Discovery of Therapeutic Options in Rare Bone Diseases. Front Endocrinol (Lausanne), 11, 405. doi:10.3389/fendo.2020.00405

29. Ralston, S. H., & Uitterlinden, A. G. (2010). Genetics of osteoporosis. Endocr Rev, 31(5), 629–662. doi:10.1210/er.2009-0044

30. Schwebach, C. L., Kudryashova, E., Zheng, W., Orchard, M., Smith, H., Runyan, L. A., … Kudryashov, D. S. (2020). Osteogenesis imperfecta mutations in plastin 3 lead to impaired calcium regulation of actin bundling. Bone Res, 8, 21. doi:10.1038/s41413-020-0095-2

31. Valimaki, V. V., Makitie, O., Pereira, R., Laine, C., Wesseling-Perry, K., Maatta, J., … Valimaki, M. J. (2017). Teriparatide Treatment in Patients With WNT1 or PLS3 Mutation-Related Early-Onset Osteoporosis: A Pilot Study. J Clin Endocrinol Metab, 102(2), 535–544. doi:10.1210/jc.2016-2423

32. van Dijk, F. S., Zillikens, M. C., Micha, D., Riessland, M., Marcelis, C. L., de Die-Smulders, C. E., … Pals, G. (2013). PLS3 mutations in X-linked osteoporosis with fractures. N Engl J Med, 369(16), 1529–1536. doi:10.1056/NEJMoa1308223

33. Wang, L., Bian, X., Cheng, G., Zhao, P., Xiang, X., Tian, W., … Zhai, Q. (2020). A novel nonsense variant in PLS3 causes X-linked osteoporosis in a Chinese family. Ann Hum Genet, 84(1), 92–96. doi:10.1111/ahg.12344

34. Wang, L., Zhai, Q., Zhao, P., Xiang, X., Zhang, X., Tian, W., & Li, T. (2018). Functional analysis of p.Ala253_Leu254insAsn mutation in PLS3 responsible for X-linked osteoporosis. Clin Genet, 93(1), 178–181. doi:10.1111/cge.13081

35. Wolff, L., Strathmann, E. A., Müller, I., Mählich, D., Veltman, C., Niehoff, A., & Wirth, B. (2021). Plastin 3 in health and disease: a matter of balance. Cell Mol Life Sci, 78(13), 5275–5301. doi:10.1007/s00018-021-03843-5

36. Yorgan, T. A., Sari, H., Rolvien, T., Windhorst, S., Failla, A. V., Kornak, U., … Schinke, T. (2020). Mice lacking plastin-3 display a specific defect of cortical bone acquisition. Bone, 130, 115062. doi:10.1016/j.bone.2019.115062

